# TRIMER: Transcription Regulation Integrated with MEtabolic Regulation

**DOI:** 10.1101/2021.04.09.439167

**Authors:** Puhua Niu, Maria J. Soto, Byung-Jun Yoon, Edward R. Dougherty, Francis J. Alexander, Ian Blaby, Xiaoning Qian

## Abstract

Advances in bioengineering have enabled numerous bio-based commodities. Yet most traditional approaches do not extend beyond a single metabolic pathway and do not attempt to modify gene regulatory networks in order to buffer metabolic perturbations. This is despite access to near universal technologies allowing genome-scale engineering. To help overcome this limitation, we have developed a pipeline enabling analysis of Transcription Regulation Integrated with MEtabolic Regulation (TRIMER). TRIMER utilizes a Bayesian network (BN) inferred from transcriptomic data to model the transcription factor regulatory network. TRIMER then infers the probabilities of gene states that are of relevance to the metabolism of interest, and predicts metabolic fluxes resulting from deletion of transcription factors at the genome scale. Additionally, we have developed a simulation framework to mimic the TF-regulated metabolic network, capable of generating both gene expression states and metabolic fluxes, thereby providing a fair evaluation platform for benchmarking models and predictions. Here, we present this computational pipeline. We demonstrate TRIMER’s applicability to both simulated and experimental data and show that it outperforms current approaches on both data types.

## INTRODUCTION

There has been extensive research in *in silico* modeling and prediction of genome-scale metabolic reaction networks, mostly focusing on mutant strain design (1, 2, 3, 4, 5, 6, 7, 8, 9, 10, 11). However, living systems involve complex and often stochastic processes arising from interactions between different types of biomolecules. For more accurate and robust prediction of target metabolic behavior under different conditions or contexts, not only metabolic reactions, but also the integration of genetic regulatory relationships involving transcription factors (TFs) that may regulate metabolic reactions, should be appropriately modeled. Due to the increasing computational complexity when considering multiple types of biomolecules in one computational system model, there have not been many “validated” computational tools for this purpose.

Probabilistic Regulation Of Metabolism (PROM) (12) introduced conditional probabilities, inferred from annotated (transcription factor)–(target gene)-reaction interactions and microarray data analysis, to incorporate transcriptional regulation information in genome-scale metabolic network analysis, especially aiming to better model condition-specific metabolism. Consequently, PROM can be considered as one of the existing integrated transcriptional-metabolic network models. IDREAM (13), an updated version of PROM, additionally allowed modeling subtle growth defects. Recently, an algorithm called OptRAM was developed based on IDREAM for designing optimized strains for ethanol overproduction in yeast (14).

The essential idea of PROM and its extensions is to infer conditional probabilities of the form Pr(gene=`ON/OFF` | TF=`ON/OFF`) so that metabolic reactions regulated by specific genes – for example, through the specific enzymes manifested as gene-protein-reaction (GPR) rules – can be modeled dependent on either genotypic or environmental changes by adjusting the flux constraints in the flux balance analysis (FBA) for metabolic modeling. Although it is computationally desirable to simplify the TF regulatory roles by introducing TF-gene conditional probabilities estimated by local frequentist estimates based on gene expression profiles, global TF-gene dependency structures may not be well captured. The existing models are also limited in the sense that only conditional probabilities based on univariate conditions were modeled. More flexible modeling that enables predictions with more complicated condition changes, for example, multiple TF knockouts when designing mutant strains, is still lacking in the literature.

The main aim of this paper is to introduce a new flexible analysis pipeline, *TRIMER*—Transcription Regulation Integrated with MEtabolic Regulation, for integrative systems modeling of TF-regulated metabolism. Specifically, a Bayesian network, instead of TF-gene conditional probabilities, will be first inferred based on gene expression profiles. Based on the inferred Bayesian network, given a condition (for example, multiple TF knockouts), we can infer the corresponding probabilities of gene states and consequently flux predictions can be performed by corresponding *in silico* metabolic models.

In addition to the modeling and analysis pipeline in TRIMER, we have also developed a simulator that simulates the TF-regulated metabolic network, which can generate both gene expression states and metabolic fluxes from a given hybrid model. Such a simulator provides a fair performance evaluation platform to help better benchmark new model inference and flux prediction methods.

## MATERIALS AND METHODS

We first introduce the main components of TRIMER organized in two major modules – namely, the *transcription* regulation network module and the *metabolic regulation* model – that are integrated within a unified interacting framework. The proposed hybrid model enables condition-dependent transcriptomic and metabolic predictions for both wild-type and TF-knockout mutant strains, through general Bayesian network (BN) modeling of transcriptional regulations. We also provide the details of our TF knockout experiments from which the experimentally observed fluxes validate the *in silico* flux predictions made by TRIMER.

### Transcription Regulation Inference in TRIMER

#### Gene expression data preprocessing

The gene expression data need to be discretized for BN learning as in TRIMER, the *TF-Regulated gene Network (TRN)* concerns ‘ON/OFF’ states of TFs and genes in the network. In our implementation, quantile normalization is first applied to raw data. Then the threshold for a given quantile value is computed and data is binarized according to the threshold. The choice of the quantile value can be either set manually or be similarly determined as in PROM. In other words, we search for the best value based on the prediction performance of the learned BN. Based on the results of our experiments, the suggested quantile value for thresholding is in the range of [0.3,0.4].

#### BN learning

The key component of TRIMER is to model the genome-scale TF-regulated gene network (TRN) by a Bayesian network (BN) learned from discretized gene expression. This TRN is expected to capture the interactions between regulators (TFs) and target genes. For this purpose, we have integrated `bn-learn`, a Bioconductor package for Bayesian network modeling of biological networks (15). A naive way to learn a BN from available observed gene states is to search over the space of all possible directed acyclic graphs (DAGs) and identify the one which optimizes a given objective function evaluating the goodness of fit. However, the search space of BN model structures grows exponentially with the number of variables (nodes in the BN). Without restricting the BN structures, the BN learning can easily become infeasible when considering even less than a dozen variables. In our experiments, we implement two structure learning strategies, tree-based search for learning tree-based BN in a restricted family of Chow-Liu trees and greedy search for learning general BNs. After finding the desired BN structure, BN model parameters are estimated by maximum likelihood estimates (MLEs).

To further restrict the search space of BN structure learning, only experimentally confirmed interactions are considered as candidate edges in BNs. In this paper, for *Escherichia coli*, we have employed the interactions archived in RegulonDB (16). When needed, separate interaction inference and validation methods, such as `GENIE3` (17), `TIGRESS` (18), or `Inferelator` (19), can also serve as the prior knowledge to extend the search space for structure learning.

Tabu search, a modified hill-climbing optimization strategy, is implemented in `bn-learn` as the search method based on a chosen score function, for example, either Bayesian information criterion (BIC) or Akaike information criterion (AIC). In our implementation, we further explore the proposed bootstrap resampling in (20) to increase the exploration space and learn a more robust structure. Specifically, we search for high-score BN structures by bootstrapping multiple expression samples from the given total samples (simulated or from expression databases). The inferred edges present in at least N% of the learned BNs are finally included in the final structure. N is a threshold value, which is determined automatically as described in (20). Such a model averaging strategy helps to establish the significance of the edges in the final “average” structure for robustness against the potential data uncertainty and scarcity. In addition to learning general BNs by a greedy Tabu search, we have also implemented Chow-Liu-tree-based BN learning in TRIMER.

#### Gene state inference

Once we have learned a BN, we can infer all the relevant conditional probabilities *Pr*(*Gene*(*s*)|*TF* (*s*)) that regulate the genome-scale metabolic network (iAF1260 for *E. coli* for example) so that for TF-knockout mutants, the conditional probabilities *P*(*genes|TFs*) can model the effect of TF knockouts over the regulated target genes and therefore the corresponding metabolic fluxes at the genome scale. To do that in TRIMER without incurring high computational cost to exhaust all potential *P* (*genes|TFs*) for metabolism regulation, we only focus on the TF-target interaction list to determine which genes can be affected (annotated as target genes) when a TF is knocked out. Generally speaking, due to potential I-equivalent classes when learning BNs from data, determining the exact causal relationships from the learned BN structure is difficult. We again rely on the annotated TF-gene interaction list (in RegulonDB for example). The Kolmogorov-Smirnov test (21) is performed to select significantly coupled TF–target pairs in the interaction list. Then the filtered list is further pruned by removing the pairs that are d-separated in the learned BN. In cases when multiple TFs are knocked out at the same time, the list of the affected genes is the union of the target gene lists corresponding to each knockout TF in TRIMER. In addition, we only care about the probabilities that will affect metabolic reactions so that only the target genes that are associated with the metabolic reactions as described by the gene-protein-reaction (GPR) rules will be considered. Given this pruned interaction list, TRIMER infers corresponding conditional probabilities by BN inference algorithms. In TRIMER, exact inference is performed by the integrated package `gRain` (22) and approximate inference in `bn-learn` (15) can also be directly utilized for computational efficiency.

### Metabolic Flux Prediction in TRIMER

#### Construct transcriptional constraints over flux variables

Metabolism regulation in TRIMER is done by constructing constraints for corresponding metabolic reaction fluxes due to corresponding gene states based on GPR rules. From the BN learning and inference module, we derive a list of conditional probabilities associated with the corresponding metabolic reactions in the metabolic network model. Similar as in PROM, these probabilities together with the maximum fluxes estimated via flux variability analysis (FVA) (23) are used to constrain the reaction flux bounds through GPR rules. In the metabolic models, GPR rules are represented as Boolean expressions associated with corresponding reactions to describe the nonlinear relationships between genes and reactions. In TRIMER, we have implemented a general platform as in `TIGER` (7) to convert the conditional probability values into linear constraints over flux variables and integrate them with the metabolic model in `COBRA` (24) for flux prediction. The workflow of connecting the BN inference and metabolic flux prediction modules is illustrated in the flowchart shown in Figure 1.

**Figure 1.**
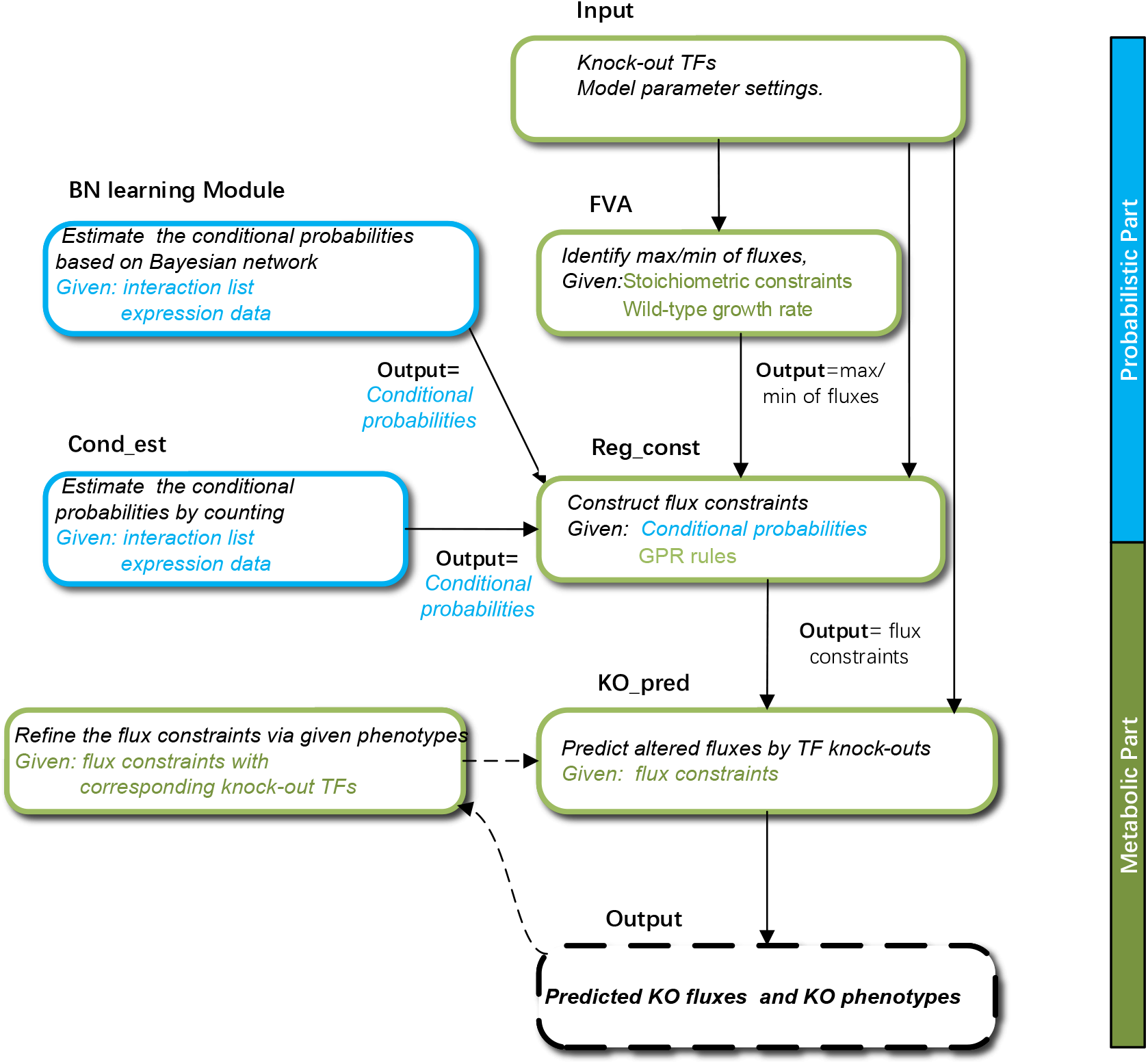
TRIMER flow-chart. The major computational modules in TRIMER and their interconnections are illustrated in the diagram.

We have adopted two ways to derive the updated reaction flux constraints according to the two ways of inferring conditional probabilities based on the learned BN. The first way is similar as the one adopted in PROM. Suppose there are M genes denoted as *G* = {*g*_1_,...,*g*_*m*_,...,*g*_*M*_} that are regulated by the corresponding TF(s). Then via the provided GPR rules in the `COBRA` model, we can find the corresponding affected reactions denoted as *R* = {*r*_1_,...,*r*_*n*_,...,*r*_*N*_}. For each *r*_*n*_∈*R*, we can find a subset of regulating genes in *G*, denoted as *G*(*r*_*n*_), based on the corresponding GPR rules. With the corresponding TF knockout mutants, the reaction flux bounds are then adjusted in the following way:

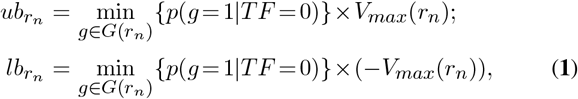

where *V*_*max*_(*r*) are estimated by FVA for reaction *r*. An example is given in the Supplemental Data to better illustrate this operation.

In TRIMER, we have also implemented a more general way for integrating both the probabilities and the GPR rules into the flux constraints, so we can obtain the joint probabilities of the states of multiple genes regulating the same reaction. The reaction flux bounds can be set by directly multiplying the maximum flux with the sum of all probabilities with the corresponding gene states that affect the corresponding reaction according to the GPR rules:

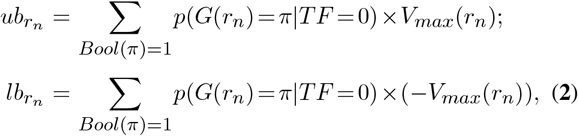

where *Bool*(*π*)=1 denotes that the corresponding GPR rules between the genes and the reaction are satisfied with *π* representing the corresponding states of genes. One illustrative example is given in the Supplemental Data.

In the remaining content, we use TRIMER-C to denote the TRIMER implementations including the flux constraints computed in the first way and TRIMER-B for the second way.

#### Data structure for metabolic reaction network

TRIMER adopts a data structure organized in a similar way as that in the `TIGER` package (7) to represent the TF-regulated metabolic reaction network. In this data structure, constraints, lower/upper bounds, variable types of the reaction flux variables provided in the model files from the `COBRA` toolbox, together with the corresponding information for additional variables are represented and stored in a unified framework. As shown in Figure 2, in the data structure representation, fields `obj, varnames, vartype, lb`, and `ub` correspond to the coefficient vector used in the objective function of the corresponding metabolic network model formulations, such as FBA and ROOM; descriptive names of involved variables; variable types; and lower/upper bounds. Fields `A, b, ctype` store all the information about constraints over variables. Stoichiometry constraints 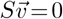 for flux variables *v* and all the other additional linear constraints over variables in the data structure specified by users are collected into matrix *A* and vector *b* and represented as a single expression 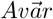 *op b*, where 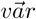 denotes all the variables included in TRIMER and *op* is a vector constituting {′>′,′<′,′=′} stored in the field `ctype`. In TRIMER, build-in functions are implemented to provide a standardized way to build the data structure mentioned above. For example, if we want to add three additional vectors 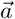, 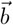 and 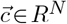 with constraints, 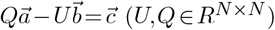 to a data structure *trimer*, it can be simply achieved by the following command lines:

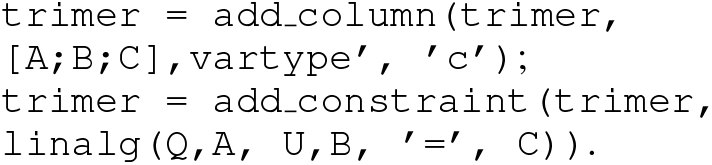

**Figure 2.**
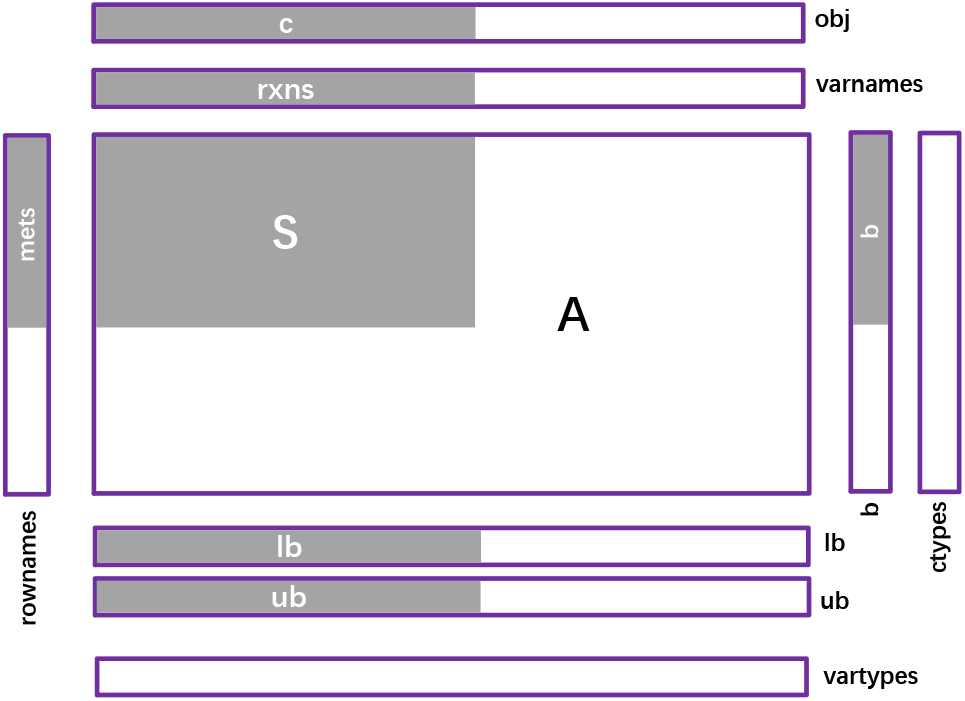
Metabolic models represented as Matlab data structures: Boxes indicate size and orientation of the fields. Black text denotes the corresponding field names. Gray areas contain data from the metabolic model, with white text indicating the relevant field names.

#### Metabolic flux prediction

Flux Balance Analysis (FBA) has been widely adopted for steady-state metabolic flux analyses by assuming the balance of production and consumption fluxes of metabolic network models (1, 9, 25). For wild-type microbial species, the corresponding steady-state fluxes can be computed by maximizing the biomass growth rate as detailed in the Supplemental Data. For modeling metabolic fluxes of mutant strains, different formulations, including minimization of metabolic adjustment (MOMA) (4) regulatory on/off minimization (ROOM) (6), OptKnock (5) and other extensions (8, 10), have been developed.

In TRIMER, we have implemented two variations of the FBA formulations for metabolic flux prediction, in addition to the standard FBA formulation with biomass as the objective function as detailed in the Supplemental Data. When predicting corresponding reaction fluxes of knockout mutants for all these formulations, let 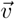,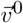*ub,lb∈ R*^*m*^ and *I*, denote the flux variables, wild-type optimal flux vector (the fluxes obtained by performing the standard FBA with the initial flux bounds given by the `COBRA` model), flux upper bounds and lower bounds, as well as the set of reactions affected by the corresponding TF knockout(s). For each affected reaction, the reaction flux bounds are modified as described previously. With that, the optimization formulation for mutants with the biomass objective and slack variables allowing violating flux bound constraints, denoted as sFBA, is as follows:

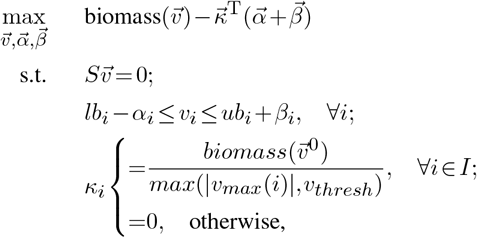

where *v*_*i*_, 1≤*i≤M* denotes the flux value for the *i*th metabolic reaction of the total *M* reactions in the metabolic network and *S* a *M × N* stoichiometric matrix with *N* metabolites in total. The biomass production flux: biomass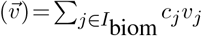 is based on the annotated set of reaction indices, *I*_biom_, involving the metabolite precursors that contribute to the biomass production in FBA with the corresponding given weights *c*_*j*_ (4). Each reaction flux is bounded by *lb*_*i*_ and *ub*_*i*_’s. Here, 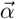 and 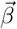 can be considered as slack variables and 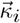 is a coefficient vector controlling which reactions are allowed to exceed the upper/lower bounds and the penalty for exceeding the bounds.

We have also implemented ROOM (6), which is believed to better model mutant strains. In the ROOM formulation, the objective is to minimize the number of reactions with significant changes from the wild-type fluxes 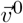. TRIMER solves the following optimization problem:

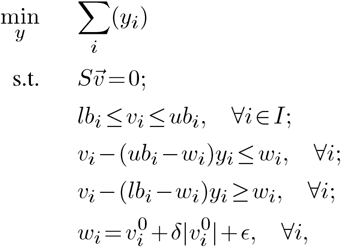

where *δ* and *ϵ* are two hyperparameters used in the original ROOM formulation to define the allowed flux changes from the wild type fluxes 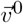.

Following `TIGER` (7), TRIMER builds a customized Matlab CMPI (Common Mathematical Programming Interface) for metabolic flux prediction based on the data structure detailed above. This CMPI defines a consistent structure for mathematical programming solvers, including `CPLEX` and `GLPK`.

### Datasets and Software Packages

TRIMER integrates several existing packages. For the BN learning and inference module, `bn-learn` (15) and `gRain` (22) are adopted for Bayesian network learning and inference respectively. For the metabolic flux prediction module, TRIMER supports `CPLEX` and `GLPK` as solvers for the three aforementioned FBA formulations. Besides, TRIMER is also compatible with the CMPI module in that `TIGER` package (7) to interface with the corresponding FBA solvers.

#### EcoMAC dataset

We have focused on the analyses with *E. coli* in TRIMER. To infer the TF regulation network and determine the ‘ON/OFF’ gene states, quantile normalization is performed over the archived microarray data in EcoMAC (26) as described previously.

#### TF-gene interaction annotations for E. coli

We have used the interaction set previously reported (26). These data comprise all archived interactions in RegulonDB v8.1 (16) that were experimentally validated to support the existence of regulatory interactions, and total 3,704 pairs of regulatory interactions. Serving as prior knowledge, those interaction pairs helped to learn the BN from microarray data and derive the TF-target list for metabolism regulation as detailed previously.

#### Metabolic model

In general, TRIMER can take any metabolic model in the `COBRA` format based on the organism under study. We have used the iAF1260 model for *E. coli* from the `COBRA` toolbox (24) throughout all the current experiments as the lab experimental data are collected from *E. coli* wild-type strains and knockout mutants.

#### GPR rules

In `COBRA` (24), the GPR rules are provided for most of the metabolic reactions, including iAF1260. TRIMER takes these GPR rules from `COBRA` directly.

### Experimental Data Collection

#### E. coli mutants and validation

Strains deleted for genes encoding transcription factors used in this study were obtained from the Keio collection of *E. coli* mutant library. All comparisons were made to BW25113, the parent strain of the collection. Mutants were validated with internal gene-specific primers by colony PCR.

#### Kovac’s Assay for indole quantification

The amount of indole produced by each mutant of interest was quantified by Kovac’s assay as described in (27). Briefly, total indole concentrations were determined by growing strains at 37°C overnight in LB or M9 minimal media and normalized to an OD600 of 0.3 the following morning. 60 ul of Kovac’s reagent (comprised of 150 ml isoamyl alcohol (IAA), 50 ml concentrated hydrochloric acid (HCl) and 10 g of para-dimethylaminobenzaldehyde (DMAB)) was added per 200 ul of normalized culture and incubated for 2 minutes. 10 ul were subsequently removed and added to 200 ul of an HCl-IAA solution, and the absorbance measured at 540 nm. Indole concentrations were then calculated using an indole standard curve prepared in the same manner as described above.

## RESULTS AND DISCUSSION

### TRIMER: Transcription Regulation Integrated with MEtabolic Regulation

We provide a brief overview of our proposed hybrid TF-regulated metabolic network model, *TRIMER*: TRIMER differs from the existing methods in the way of systematic prediction of effective intervention strategies when applied to the transcription regulatory network for regulation of metabolism. Specifically, TRIMER is based on a Bayesian Network (BN) for learning transcription regulation from gene expression data. Instead of utilizing simple conditional probabilities of the form *Pr*(*gene* = on/off|*TF* = on/off) as in PROM (12), the BN can be used to determine a probabilistic inference of the effect of alterations (e.g., gene deletions or modulated expression levels) of multiple TFs (or genes). While the framework presented is independent of the nature of TF engineering, we focus herein on gene deletions (i.e. knockouts (KO)). Furthermore, BN modeling enables intuitive incorporation of prior knowledge (e.g., pathways or pairwise regulatory relationship between genes) for learning the TRN (such as learning its topology). There exist efficient tools for learning BNs from data, which could be utilized and adapted for our purpose.

In TRIMER, a BN is trained from the gene expression data to model the joint distribution for all the relevant TFs and genes, where the resulting BN can be subsequently used to infer the steady-state conditional probabilities of the form *Pr*(*Genes|TFs*) – i.e., the probability of gene states given the states of TFs of interest. For example, we can use the BN to estimate the probability that a target gene known to regulate a specific metabolic pathway is induced given that expression of one or more TFs is abolished by gene deletion. The estimated probabilities can be used to constrain the metabolic reaction fluxes of interest, based on which the flux changes of selected metabolites resulting from the genetic alteration (e.g., TF gene deletion) can be predicted via flux balance analysis (FBA). The gene-protein-reaction (GPR) rules, which inform us of the respective metabolic pathways regulated by different genes, are used to link the translation regulation modeled by the BN with the metabolic regulation simulated by FBA.

TRIMER, which jointly models transcription regulation and metabolic regulation via the aforedescribed hybrid approach, allows us to assess the efficacy of potential TF engineering strategies and identify the optimal strategy for modulating the metabolic fluxes of interest. The desirability of a given genetic alteration can be assessed *in silico* using TRIMER, which can be validated through actual TF deletion and screening experiments in the laboratory.

Figure 3 provides a high-level overview of TRIMER, illustrating its main workflow. As shown in this diagram, TRIMER consists of two main modules: (1) the BN module for modeling and inference of transcription regulation and (2) the metabolic flux prediction module for estimating the impact of alterations in the TRN on the metabolic outcomes. The two modules are linked to each other by the GPR rules.

**Figure 3.**
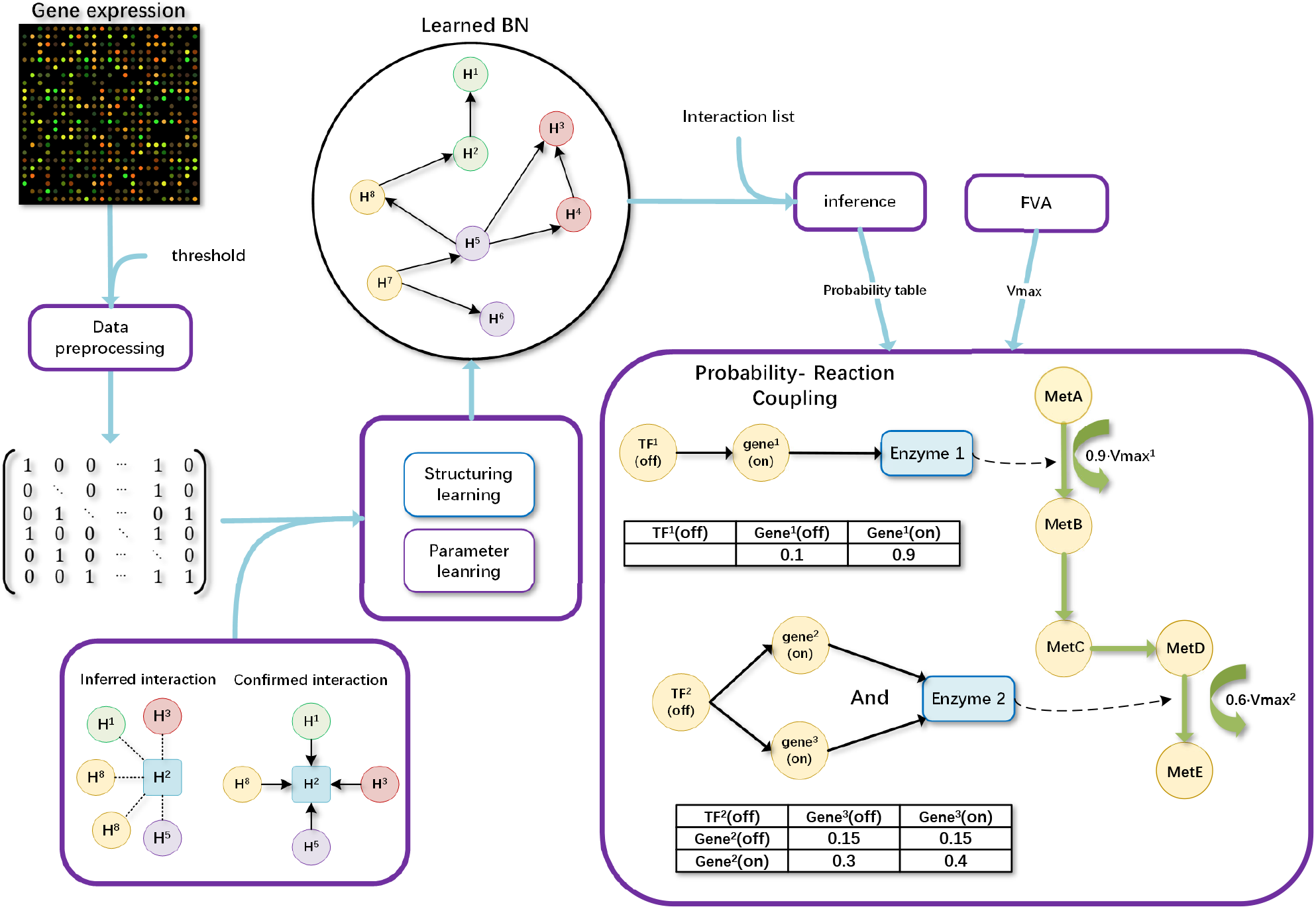
Illustrative overview of TRIMER. Gene expression data are used to infer the Bayesian network (BN) modeling the transcriptional regulations. The impact of transcription factor knock-out on downstream target genes that affect metabolic pathways are inferred using the BN. The estimated probability that a given target gene being turned on modulates the constraints in the flux variability analysis (FVA), resulting in probabilistic metabolic predictions.

Detailed description of each component can be found in the MATERIALS AND METHODS section. Furthermore, Figure 1 depicts the overall workflow in TRIMER, including the interconnections among the computational modules that comprise TRIMER.

### Simulation of *E. coli* Transcription Regulatory Network Pipeline for simulating integrated transcription and metabolic regulations

The microarray gene expression data from EcoMAC (26) are used to identify the potential target genes of 12 transcription factors studied in the aforementioned TF knockout experiments. In the current simulation experiments, we have taken these TFs and the selected target genes as the backbone nodes, in which there are 12 TFs and 32 target genes (i.e. the combined regulons). We first simulate a BN as the ground-truth TF regulation network, in which there are 137 edges in total. The interactions corresponding to these edges in this ground-truth BN are initialized with the following restrictions: 1) only the nodes corresponding to the TFs can serve as parent nodes of the nodes corresponding to other TFs or target genes; 2) the maximum number of edges between one TF parent node and all the other TF nodes is restricted to be half of the total number of TFs, and the maximum number of edges between one TF parent node and all the other target gene nodes is restricted to be half of the total number of target genes in the simulating TF regulation network; 3) the edges are randomly generated between every valid pair of nodes with the corresponding values of conditioning probability table (CPD) for each node being initialized randomly according to the uniform distribution Unif(0,1).

The gene expression data are first generated by sampling based on the simulated BN. Ten sample sets of 2000 binary gene expression profiles are drawn via the forward sampling procedure on the simulated BN. For each sample set of 2000 generated samples, five subsets of 100, 200, 400, 800, and 1600 samples are randomly selected to construct the corresponding training sets for performance evaluation. In this way, 50 datasets in total with sizes ranging from 100 to 1600 are obtained.

On the other hand, from the simulated BN, we can infer the probability of the corresponding gene states for different TF deletions as well as the wild-type. Based on the inferred probabilities and the gene-reaction relationships, the flux constraints in FBA can be adjusted to predict corresponding reaction fluxes of TF knockout mutants and the wild-type. For both the simulating ground-truth BN and the inferred BNs based on the simulated expression data, the corresponding metabolic fluxes can be simulated based on this procedure for performance evaluation. Note that all of our simulation experiments are based on the *E. coli* iAF1260 metabolic network model for FBA.

### BN structure inference based on simulated gene expression data

Given the simulated gene expression data, the first task is to learn the BN structure that best fits the given data for performance evaluation of discovering the regulatory interaction between TFs and target genes. In our experiments, we have used score-based structure learning methods for this task, where the quality of the learned BN structure is measured by the Bayesian Information Criterion (BIC) score. We have tested two BN structures: Chow-Liu tree search algorithm for identifying the global optimal tree-based BN structure and Tabu search algorithm for more general BN structure learning. Tabu search only finds the local optimal structure. In order to guarantee the quality of the predicted solutions in our experiments, the Tabu length is set to be 100 where the best structural changes in every 100 iterations are iteratively updated as a reference for future search.

Once the BN structure is inferred based on the expression data, the corresponding conditional probabilities capturing the regulation relationships can be inferred by maximum likelihood estimates (MLEs). Finally, the corresponding probabilities of gene states given TF states can be inferred by the network message passing algorithm. It should be noted that the original PROM estimates the conditional probabilities of gene states given TF states by MLE (relative frequencies) directly based on the expression data, while the authors stated that they only adjust FBA constraints by investigating only the “experimentally verified” TF-gene pairs. In our experiments, the underlying dependency between pairs of nodes in the simulating ground-truth BN is considered as the actual TF-gene pairs for PROM, to some extent in favor of PROM since it does not learn the regulation network structure.

In Table 1, we have shown the average numbers of false positive and false negative edges in the inferred general BN models by Tabu search from different numbers of training gene expression profiles. With the increasing number of training gene expression profiles, it is clear that the BN learning module in TRIMER can derive the BN model closer to the ground-truth network that is used to simulate the expression profiles as expected. Figure 4 shows the exemplar BN models learned with different numbers of training expression profiles. We have also checked the structure learning performance when we infer Chow-Liu tree based BN models, whose results are provided in Table 2. As expected, due to the imposed constraints to allow optimal search for the tree-based BN models, the true negatives are much larger compared to the results in Table 1. Nevertheless, both false positives and false negatives decrease with the increasing number of training expression profiles.

**Table 1.** False negative/positives for learned BN structures by Tabu search

| Training dataset size | 100 | 200 | 400 | 800 | 1600 |
| --- | --- | --- | --- | --- | --- |
| False positive (avg ± std) | 39.2 ± 3.7 | 23.6 ± 6.0 | 17.0 ± 3.2 | 11.8 ± 2.9 | 10.4 ± 2.5 |
| False negative (avg ± std) | 55.8 ± 3.8 | 36.6 ± 5.4 | 24.4 ± 2.2 | 15.6 ± 2.5 | 13.8 ± 2.3 |

<sup>1</sup> The total number of edges in the ground-truth simulated Bayesian network is 137.

**Figure 4.**
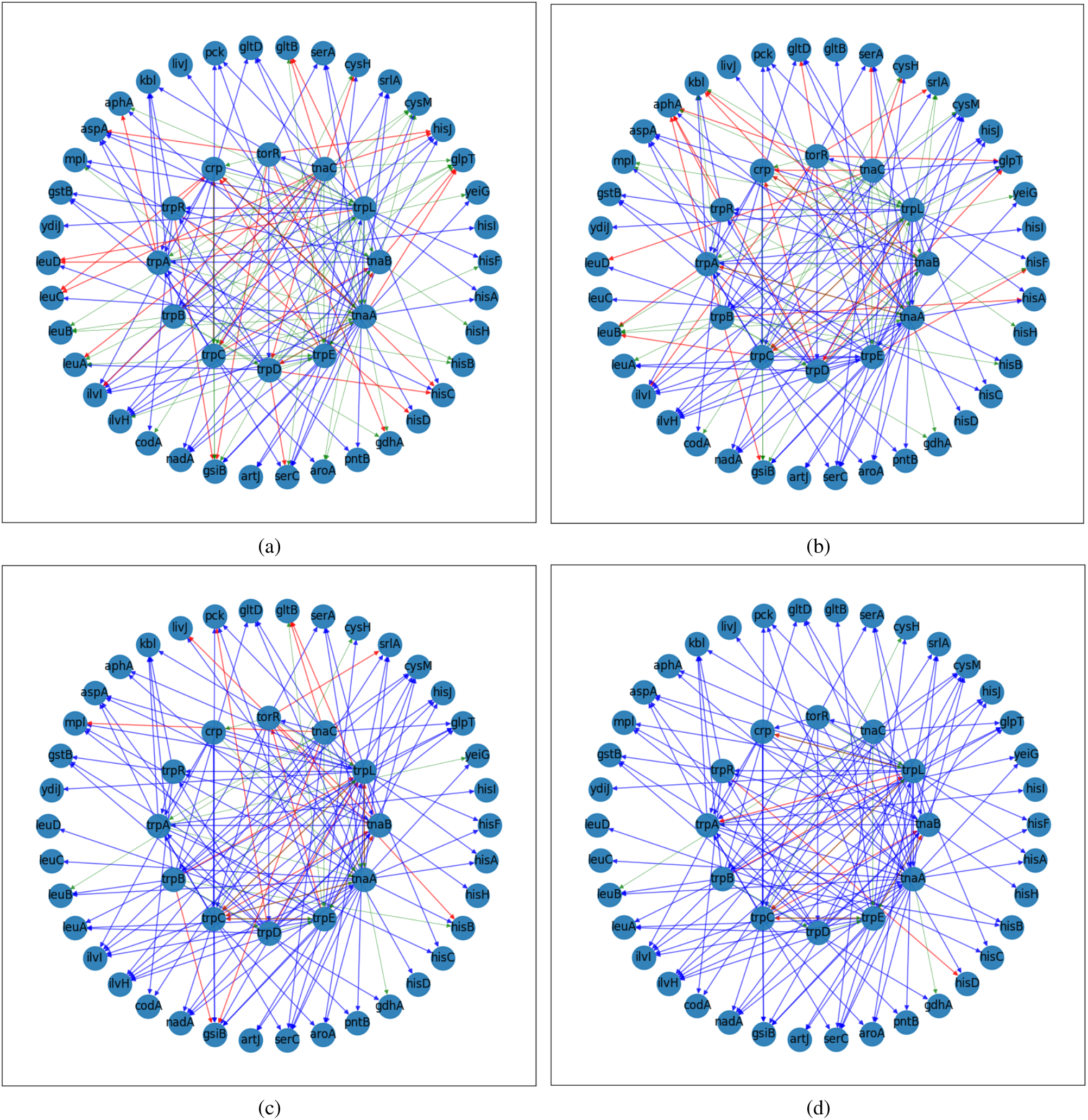
Examples of learned BNs from (a) 100, (b) 200, (c) 800, and (d) 1600 simulated expression profiles. The blue edges denote the accurately learned edges, the red edges are false positives, and the green edges are false negatives.

**Table 2.** False negative/positives for learned tree-based BN structures

| Training dataset size | 100 | 200 | 400 | 800 | 1600 |
| --- | --- | --- | --- | --- | --- |
| False positive (avg± std) | 21.8± 3.4 | 16.6± 3.1 | 13.5± 2.1 | 10.1±1.0 | 9.4±1.2 |
| False negative (avg± std) | 107.8± 3.4 | 102.6± 3.1 | 99.5± 2.1 | 96.1± 1.0 | 95.4± 1.2 |

<sup>1</sup> The total number of edges in the ground-truth simulated Bayesian network is 137.

### Evaluation of flux prediction using TRIMER based on the inferred network

In this section, we compare the flux prediction results by TRIMER-C with Chow-Liu tree (tree-TRIMER) and general BN structure (BN-TRIMER) to the results by PROM, based on simulated gene expression data. Note that we focused on applying the flux constraints based on (**1**) (TRIMER-C) for fair performance comparison with PROM. We computed the correlation between the simulated biomass and indole fluxes based on the ground-truth network model and the predicted biomass and indole fluxes based on the inferred networks of both wild-type and the mutant strains deleted for TFs in the regulation network. For 10 simulated datasets of the same number of gene expression profiles, the average correlation and its standard deviation (std) are computed. Tables 3 and 4 summarize the performance comparison of TRIMER and PROM for biomass and indole flux prediction respectively, with different numbers of simulated gene expression data.

**Table 3.** Biomass flux prediction comparison between TRIMER and PROM with different numbers of training expression profiles

| average/std | 100 | 200 | 400 | 800 | 1600 |
| --- | --- | --- | --- | --- | --- |
| BN-TRIMER | 0.631±0.199 | 0.847±0.181 | 0.936±0.062 | 0.978±0.015 | 0.986±0.005 |
| tree-TRIMER | 0.432±0.276 | 0.565±0.170 | 0.661±0.117 | 0.605±0.145 | 0.599±0.178 |
| PROM | 0.125±0.275 | 0.488±0.362 | 0.775±0.133 | 0.659±0.145 | 0.788±0.163 |

**Table 4.** Indole flux prediction comparison between TRIMER and PROM with different numbers of training expression profiles

| average/std | 100 | 200 | 400 | 800 | 1600 |
| --- | --- | --- | --- | --- | --- |
| BN-TRIMER | 0.873±0.086 | 0.949±0.071 | 0.979±0.019 | 0.921±0.221 | 0.917±0.248 |
| tree-TRIMER | 0.863±0.053 | 0.902±0.030 | 0.914±0.030 | 0.911±0.034 | 0.913±0.035 |
| PROM | 0.382±0.253 | 0.303±0.253 | 0.485±0.227 | 0.699±0.155 | 0.786±0.232 |

As shown in Table 3, from simulated expression data, BN-TRIMER consistently gives the closest biomass flux prediction to the simulated fluxes based on the ground-truth model. It is clear that with more expression data, the predicted fluxes can get better and vary less with different simulated expression data. With small training expression data, PROM’s flux prediction can have quite weak correlation while with increasing size of expression data, the prediction can improve. For tree-TRIMER, as the model class deviates from the ground-truth model, the prediction performance saturates when the number of training expression profiles is 400. On the other hand, with small training sets, tree-TRIMER performs better than PROM.

Table 4 provides a comparison for the indole flux predictions. Note that the ground-truth BN models are simulated based on the core subnetwork centering around indole-related reactions. We observe that both versions of TRIMER-C have better indole flux prediction performance, especially with small training data, compared to the results in Table 4. The tree-TRIMER shows much better performance, which suggests that good prior knowledge on what to model for the TF regulation network may significantly enhance flux predictions. On the other hand, unlike TRIMER, PROM only models local dependency instead of global dependency, and its indole and biomass flux prediction performances are similar. In Figures 5 and 6, we provide the bar plots based on the comparison previously discussed.

**Figure 5.**
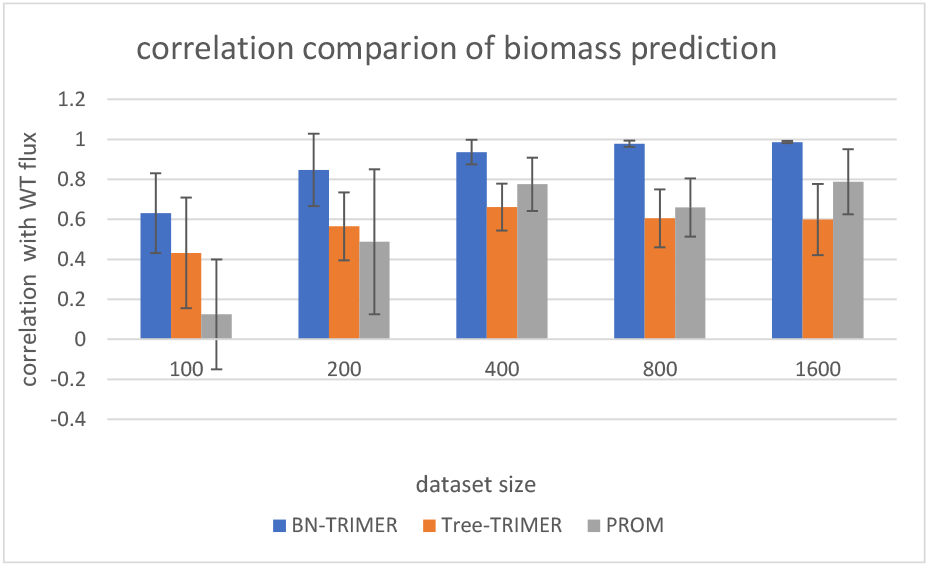
Biomass flux prediction comparison between TRIMER and PROM

**Figure 6.**
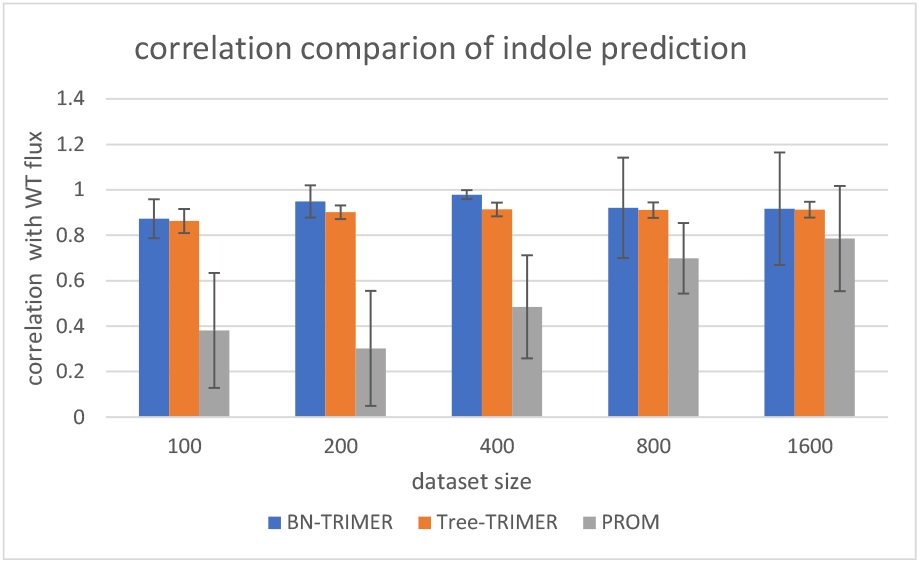
Indole flux prediction comparison between TRIMER and PROM

### Experimental validation of metabolic flux predictions made by TRIMER

To further demonstrate the utility of TRIMER in *in silico* metabolic flux prediction for TF knockout mutants, we have taken the archived microarray gene expression data and the experimentally verified TF-gene interactions in EcoMAC (26) to infer the corresponding Bayesian network for modeling the TF regulation network using the general BN inference module of TRIMER. Based on the inferred BN, the conditional probabilities of corresponding gene states when given TF knockouts were computed. Taking these inferred probabilities, the metabolic network flux prediction module in TRIMER was run to predict biomass and indole fluxes for corresponding TF knockout mutants.

We first compared the *in silico* flux predictions by TRIMER and PROM with the experimental measurements from the knockout experiments in (28), for which the prediction results by PROM were reported in (12). For the biomass objective, we took `Ec_biomass_iAF1260_core_59p81M` in iAF1260 as done in PROM. Three FBA formulations, standard FBA, sFBA, and ROOM in TRIMER were implemented. In our experiments, the parameters δ and ε in the ROOM formulation were set to be 0.05 and 0.001. Table 5 provides the comparison of the experimental and predicted fluxes by TRIMER-C, TRIMER-B, and PROM for different TF knockout mutants as well as the overall Pearson’s correlation coefficients. TRIMER-B consistently achieved the highest correlation with the experimental results for three FBA formulations, among which TRIMER-B with sFBA obtained the highest correlation with the experimental results among all the model choices.

**Table 5.** Predicted biomass flux comparison between TRIMER and PROM for the knockout experiments in (28).

| TF KO | Actual | TRIMER-C |  |  | TRIMER-B |  |  | PROM |  |  |
| --- | --- | --- | --- | --- | --- | --- | --- | --- | --- | --- |
|  |  | FBA | sFBA | ROOM | FBA | sFBA | ROOM | FBA | PROM | ROOM |
| WT +O2 | 0.71 | 0.7077 | 0.7077 | 0.7077 | 0.7077 | 0.7077 | 0.7077 | 0.7077 | 0.7077 | 0.7077 |
| arcA+O2 | 0.6900 | 0.1006 | 0.5828 | 0.0977 | 0.2784 | 0.6091 | 0.2538 | 0.1967 | 0.2722 | 0.0418 |
| fnr +O2 | 0.6300 | 0.4109 | 0.5227 | 0.3557 | 0.3852 | 0.5363 | 0.3852 | 0.3994 | 0.5256 | 0.3557 |
| arcA fnr +O2 | 0.6500 | 0.1006 | 0.4109 | 0.0863 | 0.2784 | 0.5363 | 0.2653 | 0.1967 | 0.2722 | 0.0399 |
| appY +O2 | 0.6400 | 0.7077 | 0.7077 | 0.6714 | 0.7077 | 0.7077 | 0.6714 | 0.7077 | 0.7077 | 0.6714 |
| oxyR +O2 | 0.6400 | 0.7077 | 0.7077 | 0.6714 | 0.7077 | 0.7077 | 0.6714 | 0.7077 | 0.7077 | 0.6714 |
| soxS +O2 | 0.7200 | 0.6524 | 0.7063 | 0.6524 | 0.6476 | 0.7062 | 0.6476 | 0.6493 | 0.7073 | 0.6493 |
| WT -O2 | 0.49 | 0.4914 | 0.4914 | 0.4914 | 0.4914 | 0.4914 | 0.4914 | 0.4914 | 0.4914 | 0.4914 |
| arcA -O2 | 0.3800 | 0.0193 | 0.0193 | 0.0097 | 0.0535 | 0.0535 | 0.0482 | 0.0378 | 0.0378 | 0.0278 |
| fnr -O2 | 0.4100 | 0.2853 | 0.3629 | 0.2853 | 0.2674 | 0.3724 | 0.2674 | 0.1421 | 0.2773 | 0.1224 |
| arcA fnr -O2 | 0.3000 | 0.0193 | 0.0193 | 0.0088 | 0.0535 | 0.0535 | 0.0484 | 0.0378 | 0.0378 | 0.0373 |
| appY -O2 | 0.4800 | 0.4914 | 0.4914 | 0.4658 | 0.4914 | 0.4914 | 0.4658 | 0.4914 | 0.4914 | 0.4658 |
| oxyR -O2 | 0.4800 | 0.4914 | 0.4914 | 0.4658 | 0.4914 | 0.4914 | 0.4658 | 0.4914 | 0.4914 | 0.4658 |
| soxS -O2 | 0.4600 | 0.4529 | 0.4914 | 0.4529 | 0.4496 | 0.4914 | 0.4496 | 0.4508 | 0.4902 | 0.4508 |
| correlation | - | 0.5063 | 0.8377 | 0.5019 | **0.6557** | **0.8843** | **0.6537** | 0.6086 | 0.6793 | 0.4767 |

<sup>1</sup> In the FBA formulations, substrate (glucose) and oxygen uptake rates for aerobic conditions are set to be 8.5 and 14.6 mmol/gDCW/hr, respectively. They are set to 0 and 21.2 mmol/gDCW/hr for anaerobic conditions.
<sup>2</sup> The optimization is by the CPLEX solver.

We further validated the predicted fluxes by TRIMER with our experimentally-generated data from TF-knockout experiments for indole production as described previously. We took `TRPS3` in iAF1260 for indole flux prediction. Table 6 provides the comparison of the experimental and predicted fluxes by TRIMER-C, TRIMER-B and PROM for different TF knockout mutants grown in M9 minimal media and the overall Pearson’s correlation coefficients between experimental and predicted fluxes. In this set of experiments, TRIMER again has achieved consistently better correlation with the experimental results. The prediction performance of TRIMER-B is better with the sFBA and ROOM formulations. It should be also noted that TRIMER with the ROOM formulation has achieved the highest correlation values, which are significantly better than the other FBA formulations for both TRIMER and PROM. The overall superiority of TRIMER over PROM is due to the effective modeling of the global dependency in TF regulations through the BN learning and inference, in contrast to using simple conditional probability estimates adopted in PROM.

**Table 6.**
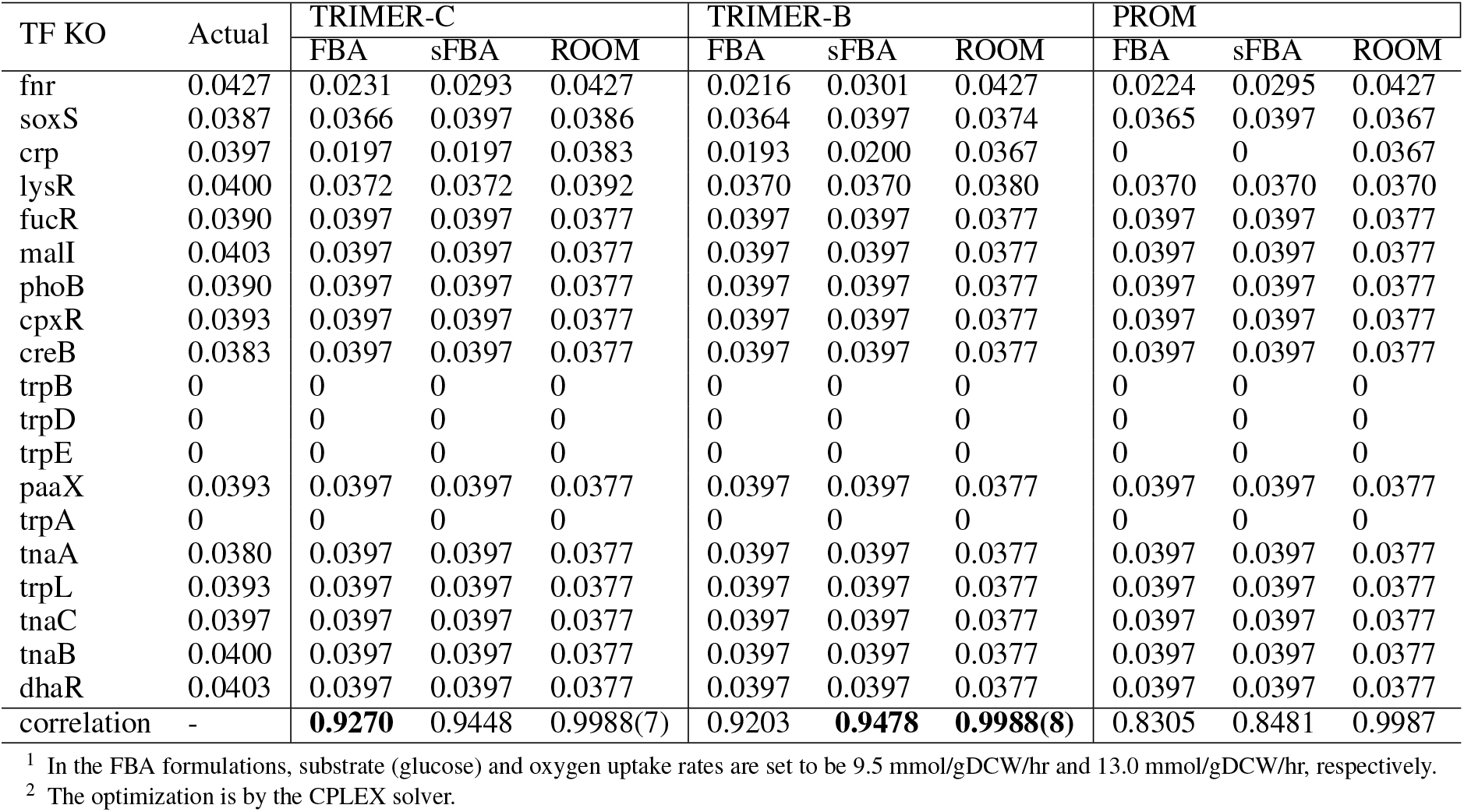
Predicted indole flux comparison between TRIMER and PROM for our TF knockout (KO) experiments in M9 minimal media.

## Supporting information

Indole concentration tables

## DATA AVAILABILITY

### TRIMER: Source Code and Instructions

The developed package TRIMER is available online in an open-source GitHub repository (29). All the implemented functions in TRIMER are documented using Matlab’s help function. Code for the reported experiments in this paper can also be found in the GitHub repository.

### Experimental Data

Experimental data for total indole concentrations of the TF-knockout deletants and the parental strain in LB and M9 media are provided as Supplementary Tables 1 and 2, respectively.

## ACKNOWLEDGEMENTS

PN and XQ are partially supported by the National Science Foundation under Grants CCF-1553281.

## SUPPLEMENTAL DATA

We here provide the brief review on the basics of flux balance analysis (FBA) (9, 25) and probabilistic regulatory network modeling including both simplistic conditional probability models as adopted in Probabilistic Regulation Of Metabolism (PROM) (12) and more general Bayesian network (BN) modeling (30). We also present some examples of potential conditional probabilities inferred from the learned BNs. Finally, we introduce, with additional experimental results, the phenotype prediction functions of TRIMER with integrated modules from `TIGER` (7).

### Flux Balance Analysis and metabolic engineering for mutant strain design

Since it has been proposed in (1, 9, 25), Flux Balance Analysis (FBA), as a simplified network analysis model for metabolic flux analysis, has been widely adopted for steady-state flux analyses by assuming the balance of production and consumption fluxes of metabolic network models. Mathematically, with the prior stoichiometry knowledge, FBA assumes that the weighted sum of network fluxes, denoted by the vector 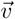, based on stoichiometric coefficients *S* is 0: 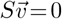. Such a steady-state flux analysis is performed by assuming that the corresponding wild-type microbial species always optimizes for its growth:

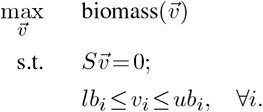

For wild-type microbial strains, a common assumption is that their steady-state flux values follow an optimal distribution that maximizes the biomass production rate. Often, the steady-state flux distribution is approximately solved as a linear programming (LP) problem to maximize the biomass production flux: 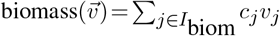 subject to the FBA stoichiometry constraints. Here, *c*_*j*_ is the corresponding given weighting coefficients and *I*_biom_ is the set of reaction indices involving the metabolite precursors that contribute to the biomass production in FBA (4).

When modeling mutant strains, the researchers found that the biomass maximization assumption for wild-type strains may not approximate the steady-state fluxes well. To achieve better agreement with experimental observations, approximation formulations of knockout metabolic fluxes undergoing a minimization of metabolic adjustment (MOMA) process (4) or by the regulatory on/off minimization (ROOM) (6) have been proposed to address the long-term post knockout metabolic flux distribution predication problem.

Existing microbial strain design formulations are mostly in the framework of FBA without considering changing conditions or contexts. They search for the knockouts to optimize the desired flux predictions by bi-level optimization formulations to make sure about the mutant survival at the same time. One of such representative methods is OptKnock (5). However, when modeling condition/context dependency in hybrid models involving transcriptional regulations, such methods are not directly applicable.

### PROM: A brief review

PROM aims to predict metabolic fluxes of the knockout mutants in transcription factor (TF)-regulated metabolic networks. Specifically, PROM is built upon the FBA framework. PROM first estimates the probability of “reaction-targeted” gene expression (ON/1 or OFF/0) given transcription factor (TF) expression PR(*gene* = 1|*TF* = 0) based on a certain set of microarray expression data using annotated TF-gene-reaction interactions. Based on that, PROM solves the following LP problem given transcription factor knockout (KO) perturbations:

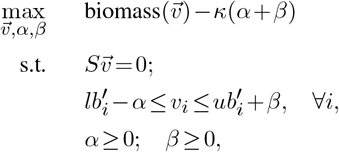

where *α* and *β* can be considered as slack variables and *lb*′_*i*_ and 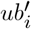 are perturbed flux bounds based on transcriptional regulations. In particular, Flux Variability Analysis (FVA) (23) is performed together with network-based metabolic behavior prediction (31) to get the minimum and maximum fluxes. The inferred conditional probabilities due to a specific TF KO will then be multiplied based on the transcriptional regulations on the corresponding metabolic reactions, for which the metabolic models from either the KBase (32) or `COBRA` toolbox (24) can be used.

PROM consists of multiple steps from microarray data analysis, flux bound manipulations, and FBA based on these steps with the aim to have their model prediction to better fit with the flux measurements at different conditions.

### Modeling transcription regulations using Bayesian Network

A Bayesian network (BN) is a probabilistic graphical model (PGM) that can be used to represent the joint probability distribution of a set of variables **X** = {*X*_1_,...,*X*_*n*_}, whose dependencies are described by a directed acyclic graph (DAG) 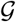. Each node in the DAG corresponds to a variable *X*_*i*_*in***X** of interest, and a directed edge *X*_*j*_→*X*_*k*_ represents the possible causal relationship between the variables *X*_*j*_ and *X*_*k*_. Following the topology of the DAG, the joint distribution of **X** can be written as a product of conditional probabilities:

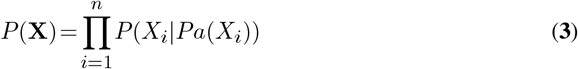

where *P*(*X*_*i*_|*Pa*(*X*_*i*_)) is the conditional distribution function of *X*_*i*_ given the set of variables *Pa*(*X*_*i*_), which denote the set of its parent nodes in 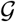. In BN, graph topology captures the complex dependencies among the variables, resulting in a compact representation of the joint probability distribution of **X** by factorizing it into a product of local probability models as in (**3**). This compact representation reduces data requirements for learning the distribution from data and also greatly enhances the computational efficiency of making probabilistic inference based on the distribution (30).

The main novelty of the hybrid models in TRIMER is to model the TF-regulated network (TRN) using the more general Bayesian network model to better capture regulatory relationships. Unlike PROM, in which TF-regulations are represented simply by inferring the maximum likelihood estimates (MLEs) of the involved conditional probabilities PR(*gene*= 1|*TF* = 0), TRIMER adopts a full-fledged BN to capture the transcription regulations. Based on the available gene expression data, we can infer the BN by first inferring the structure of the network and then estimating the parameters of the local probability model (i.e., conditional probability distributions). In this manner, TRIMER can better capture both the local and global dependencies between TFs and genes, thereby better model the TF knockout effects on metabolic fluxes. Furthermore, the BN enables the incorporation of available prior knowledge regarding TF regulations, enhancing the quality of the inferred network compared to a solely data-driven inference approach.

### Examples of inferring conditional probabilities given BN

In this subsection, we provide two examples for the two ways of applying transcriptional regulations based on BN-inferred conditional probabilities as explained in the main text.

We provide an example to illustrate the operation based on (**1**). In this example, the reaction *r* is catalyzed by two genes *A* and *B* according to the GPR rules in the metabolic model. When their regulating TF is knockout, we can obtain a probability vector: (*P*(*A* = 1|*TF* = 0)*,P* (*B* = 1|*TF* = *on*))^T^. The corresponding reaction flux upper/lower bounds for reaction *r* are set to be:

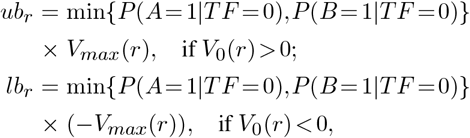

where *V*_0_(*r*) is the wild-type flux for reaction *r*.

We now give the example to illustrate the operation based on (**2**). In this example, the reaction *r* is associated with a GPR rule, (*A and B*)*orC*. The corresponding GPR rule values and three gene states are illustrated in Table A1. We can see that only four of the sixteen possible state combinations render that the GPR rule to be false. Thus, the upper or lower bounds with respect to *r* will be computed as:

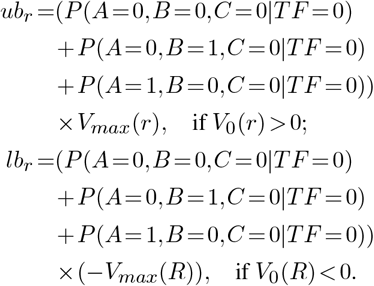

**Table A1.**
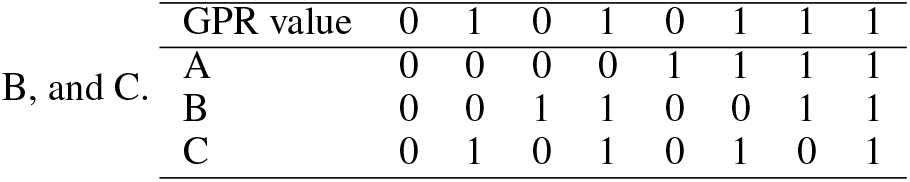
GPR rules for gene state profiles of three genes: A,

### Refine TRIMER with given phenotypes

In TRIMER, we provide a way to refine the current metabolic model given a minimum growth rate. This can help to remove or adjust regulatory bounds that over-constrain the prediction model when TFs are knocked out. These bounds can be decided by solving the following optimization problem:

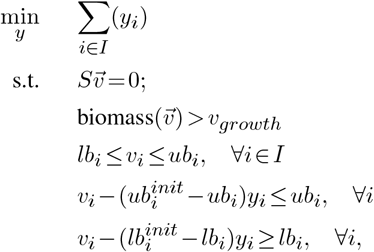

where *ub*^*init*^ and *lb*^*init*^ are the initial bounds from the original wild-type model in `COBRA` and *v*_*growth*_ is the minimum growth-rate requirement specified by the user. Suppose the threshold for a lethal KO is marked with 0.05 times the wild-type biomass flux. In the experiments of biomass prediction based on the experimental data in (28), we predict that the phenotype of arcA KO is lethal when we have sFBA with the TRIMER-C model. However, the actual phenotype of arcA KO is non-lethal according to the experimentally measured fluxes. This may indicate that some estimated conditional probabilities for constructing flux constraints are too small and some reactions affected by arcA KO are over-constrained. Via the optimization problem above, we can identify which reactions are over-constrained for TF(s) knockouts for given non-lethal phenotypes. Based on this, we can further adjust the values of conditional probabilities corresponding to these reactions to make the predicted phenotypes to better match the experimentally observed phenotypes and also, the predicted fluxes to be closer to the experimentally observed fluxes. The name abbreviations of the reactions that are over-constrained and their corresponding condition probabilities for arcA KO is shown in Table A2. It is clear that these probability values are all small and result in the predicted phenotype to be lethal. We adjust all these probabilities to be 0.9 and the predicted biomass flux becomes 0.4027, which is very close to the experimentally measured flux.

### Integrating TRIMER with TIGER

In TRIMER, we have also programmed a simulation pipeline that can simulate knockout mutant metabolic fluxes in various growth conditions by borrowing the modules in `TIGER` (7), instead of only being capable simulating aerobic and anaerobic glucose minimal medium conditions as in PROM (12). We adopt a Boolean model to simulate the feedback regulatory rules as implemented in `TIGER`. As TRIMER adopts the similar data structure as `TIGER`, which is known to be a platform to integrate `COBRA` models with these Boolean transcription regulations, TRIMER allows the user to build a hybrid model that integrates probabilistic TRN, Boolean feedback rules, and `COBRA` metabolic models into a single unified pipeline, making it possible to simulate knockout mutants in various growth conditions. We have simulated 125 growth conditions for 15 TF KOs based on the *E. coli* iAF1260 model and compared the performance of this hybrid model and that by PROM using the phenotype datasets, originally given in (12), as the ground truth. The parameter settings of growth conditions can be found in (28). For the `TIGER` part of the hybrid model to model Boolean regulations interfacting the TRN and metabolic model, we have adopted the iMC1010 Boolean network in (28).The TRIMER part of the hybrid model is the same as PROM. Figure A1 shows the results. The best performance of the hybrid model and PROM implentations are both achieved when the threshhold for lethal phenotypes is set to be 0.15 times the WT growth rate. As we can see, the predictions mainly differ in the growth condition with the growth media, 1,2-Propanediol L-Tartaric Acid, L-Tartaric Acid, and Guanine. With additional constraints introduced from the Boolean rules, many predicted phenotypes become lethal.

**Table A2.**
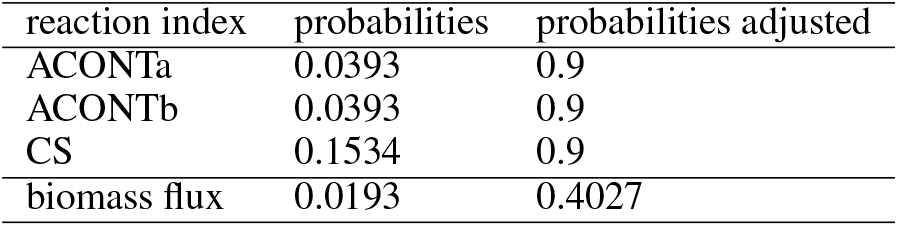
Reactions that are over-constrained with the corresponding inferred and adjusted probabilities

### TF-knockout experimental data

Total indole concentrations for the TF-knockout deletants and the parental strain used in this study are provided in the separate Supplementary Tables 1 and 2.

**Figure A1.**
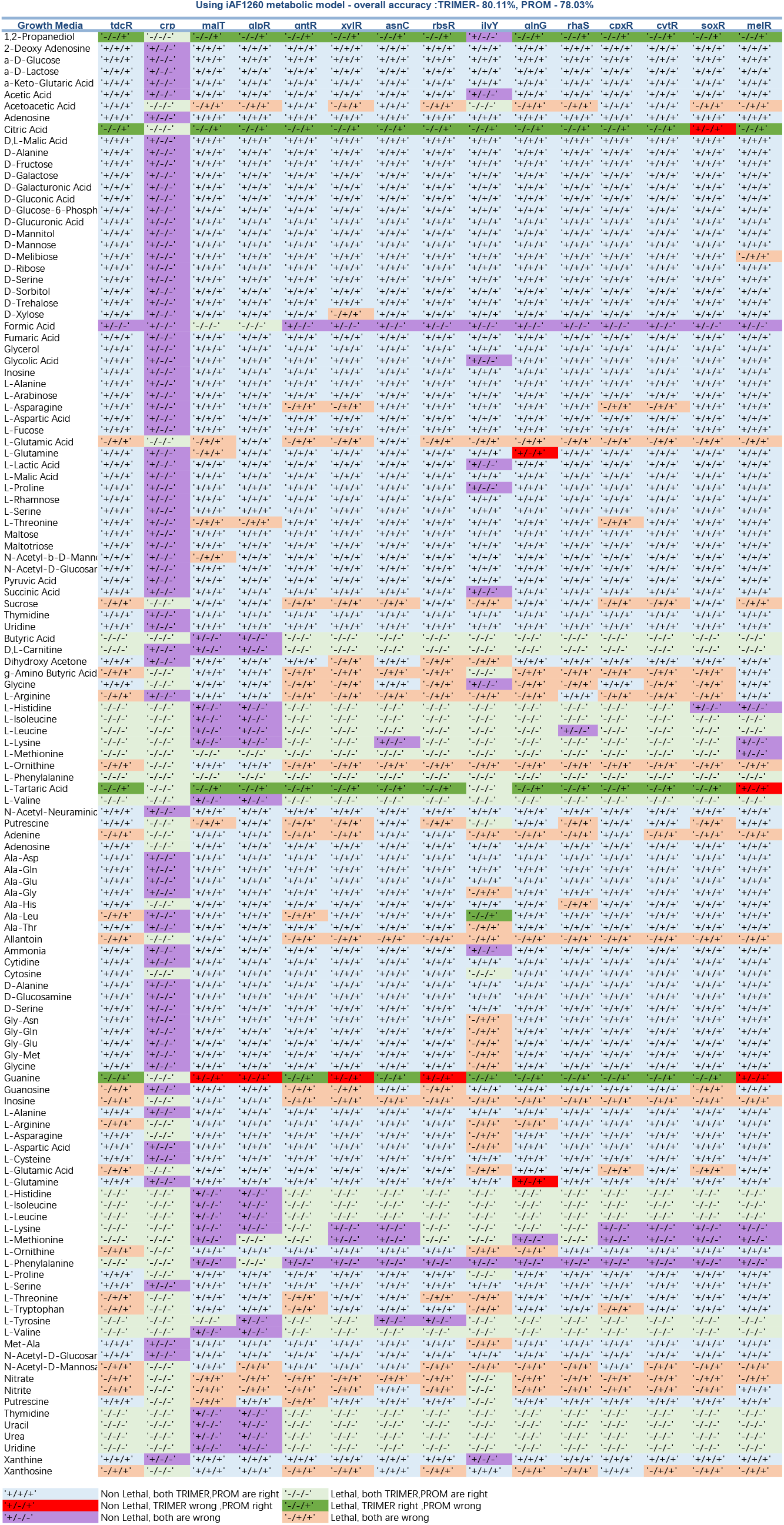
Phenotype prediction comparison between TRIMER and PROM

