## Supplementary material for "TRIMER: Transcription Regulation Integrated with MEtabolic Regulation": Indole concentration tables

**Supplementaty Table 1. Total indole concentrations of *E. coli* transcription factor deletants in LB.**

| Strain | Mean absorbance[1] | STDEV | mmols of Indole |
|---|---|---|---|
| **WT** | 0.102 | 0.006 | 0.1031 |
| *crp* | 0.043 | 0.005 | 0.0000 |
| *tnaA* | 0.045 | 0.006 | 0.0000 |
| *lldR* | 0.074 | 0.006 | 0.0492 |
| *glcC* | 0.076 | 0.009 | 0.0531 |
| *nadR* | 0.080 | 0.006 | 0.0595 |
| *relB* | 0.081 | 0.006 | 0.0614 |
| *fabR* | 0.081 | 0.003 | 0.0620 |
| *arsR* | 0.081 | 0.007 | 0.0627 |
| *acrR* | 0.082 | 0.003 | 0.0640 |
| *fhlA* | 0.085 | 0.003 | 0.0697 |
| *araC* | 0.085 | 0.005 | 0.0704 |
| *marR* | 0.086 | 0.003 | 0.0710 |
| *uxuR* | 0.087 | 0.013 | 0.0736 |
| *metR* | 0.087 | 0.009 | 0.0742 |
| *cytr* | 0.088 | 0.013 | 0.0761 |
| *dsdX* | 0.089 | 0.010 | 0.0768 |
| *allR* | 0.089 | 0.004 | 0.0781 |
| *exuR* | 0.090 | 0.007 | 0.0787 |
| *soxR* | 0.090 | 0.004 | 0.0793 |
| *gadX* | 0.091 | 0.012 | 0.0819 |
| *fnr* | 0.094 | 0.003 | 0.0870 |
| *mprA* | 0.095 | 0.010 | 0.0883 |
| *nanR* | 0.095 | 0.005 | 0.0896 |
| *stpA* | 0.096 | 0.009 | 0.0902 |
| *nhaR* | 0.096 | 0.013 | 0.0915 |
| *kdgR* | 0.097 | 0.005 | 0.0922 |
| *idnR* | 0.097 | 0.008 | 0.0922 |
| *melR* | 0.097 | 0.005 | 0.0928 |
| *ada* | 0.098 | 0.012 | 0.0941 |
| *metJ* | 0.098 | 0.015 | 0.0941 |
| *mlrA* | 0.098 | 0.007 | 0.0941 |
| *galS* | 0.098 | 0.019 | 0.0954 |
| *tyrR* | 0.099 | 0.010 | 0.0967 |
| *ilvY* | 0.099 | 0.017 | 0.0967 |
| *xapR* | 0.100 | 0.017 | 0.0979 |
| *zntR* | 0.100 | 0.012 | 0.0979 |
| *rstA* | 0.100 | 0.012 | 0.0979 |
| *nagC* | 0.100 | 0.006 | 0.0992 |
| *csiR* | 0.100 | 0.004 | 0.0992 |
| *hupA* | 0.101 | 0.010 | 0.1011 |
| *trpA* | 0.101 | 0.006 | 0.1011 |
| *leuO* | 0.101 | 0.008 | 0.1011 |
| *ebgR* | 0.102 | 0.018 | 0.1024 |

| | | | |
|---|---|---|---|
| *lacI* | 0.102 | 0.004 | 0.1031 |
| *soxS* | 0.103 | 0.017 | 0.1050 |
| *rtcR* | 0.104 | 0.007 | 0.1069 |
| *rbsR* | 0.105 | 0.011 | 0.1076 |
| *narL* | 0.105 | 0.014 | 0.1082 |
| *gntR* | 0.105 | 0.008 | 0.1082 |
| *chbR* | 0.106 | 0.021 | 0.1095 |
| *lysR* | 0.106 | 0.018 | 0.1108 |
| *glnG* | 0.106 | 0.005 | 0.1108 |
| *lrp* | 0.107 | 0.014 | 0.1127 |
| *cbl* | 0.107 | 0.011 | 0.1127 |
| *rhaS* | 0.107 | 0.010 | 0.1127 |
| *sgrR* | 0.108 | 0.010 | 0.1140 |
| *yehT* | 0.108 | 0.010 | 0.1140 |
| *envR* | 0.108 | 0.015 | 0.1146 |
| *glpR* | 0.109 | 0.001 | 0.1159 |
| *fadR* | 0.110 | 0.010 | 0.1172 |
| *xylR* | 0.110 | 0.013 | 0.1178 |
| *uidR* | 0.110 | 0.012 | 0.1184 |
| *torR* | 0.111 | 0.029 | 0.1191 |
| *oxyR* | 0.111 | 0.011 | 0.1197 |
| *rhaR* | 0.111 | 0.014 | 0.1204 |
| *rpiR* | 0.111 | 0.007 | 0.1204 |
| *appY* | 0.112 | 0.017 | 0.1210 |
| *creB* | 0.112 | 0.013 | 0.1210 |
| *hcaR* | 0.112 | 0.004 | 0.1210 |
| *slyA* | 0.112 | 0.003 | 0.1210 |
| *prpR* | 0.112 | 0.011 | 0.1217 |
| *feaR* | 0.112 | 0.010 | 0.1217 |
| *srlR* | 0.112 | 0.016 | 0.1223 |
| *dcuR* | 0.112 | 0.014 | 0.1223 |
| *argP* | 0.113 | 0.009 | 0.1242 |
| *caiF* | 0.113 | 0.028 | 0.1242 |
| *nac* | 0.114 | 0.020 | 0.1249 |
| *cynR* | 0.114 | 0.012 | 0.1249 |
| *betI* | 0.114 | 0.013 | 0.1249 |
| *aidB* | 0.114 | 0.008 | 0.1249 |
| *fis* | 0.114 | 0.016 | 0.1255 |
| *adiY* | 0.114 | 0.009 | 0.1255 |
| *trpC* | 0.114 | 0.010 | 0.1255 |
| *galR* | 0.114 | 0.003 | 0.1261 |
| *fliZ* | 0.114 | 0.009 | 0.1261 |
| *argR* | 0.115 | 0.014 | 0.1268 |
| *tdcA* | 0.115 | 0.015 | 0.1274 |
| *evgA* | 0.115 | 0.017 | 0.1274 |
| *gadE* | 0.115 | 0.009 | 0.1274 |
| *gadW* | 0.115 | 0.010 | 0.1274 |

| | | | |
|---|---|---|---|
| *gutM* | 0.116 | 0.002 | 0.1293 |
| *cdaR* | 0.116 | 0.011 | 0.1300 |
| *ycfQ* | 0.116 | 0.009 | 0.1300 |
| *arcA* | 0.117 | 0.007 | 0.1319 |
| *marA* | 0.117 | 0.007 | 0.1319 |
| *bglJ* | 0.117 | 0.004 | 0.1319 |
| *trpL* | 0.118 | 0.013 | 0.1326 |
| *treR* | 0.118 | 0.019 | 0.1332 |
| *phoP* | 0.118 | 0.021 | 0.1332 |
| *iscR* | 0.118 | 0.015 | 0.1338 |
| *paaX* | 0.118 | 0.011 | 0.1338 |
| *fur* | 0.118 | 0.008 | 0.1338 |
| *tnaC* | 0.120 | 0.015 | 0.1370 |
| *mngR* | 0.120 | 0.013 | 0.1370 |
| *cueR* | 0.121 | 0.003 | 0.1383 |
| *rob* | 0.121 | 0.016 | 0.1383 |
| *tnaB* | 0.121 | 0.012 | 0.1383 |
| *csgD* | 0.121 | 0.006 | 0.1390 |
| *asnC* | 0.121 | 0.011 | 0.1390 |
| *cadC* | 0.122 | 0.015 | 0.1402 |
| *mhpR* | 0.122 | 0.006 | 0.1402 |
| *yeiL* | 0.122 | 0.004 | 0.1402 |
| *cspA* | 0.122 | 0.007 | 0.1402 |
| *kdpE* | 0.122 | 0.006 | 0.1409 |
| *gatR* | 0.122 | 0.005 | 0.1415 |
| *bolA* | 0.124 | 0.010 | 0.1441 |
| *norR* | 0.124 | 0.009 | 0.1454 |
| *sdiA* | 0.125 | 0.014 | 0.1460 |
| *malI* | 0.125 | 0.017 | 0.1460 |
| *purR* | 0.125 | 0.008 | 0.1467 |
| *lrhA* | 0.125 | 0.010 | 0.1467 |
| *zur* | 0.126 | 0.010 | 0.1479 |
| *narP* | 0.126 | 0.015 | 0.1486 |
| *basR* | 0.126 | 0.018 | 0.1492 |
| *alaS* | 0.126 | 0.015 | 0.1492 |
| *atoC* | 0.128 | 0.022 | 0.1524 |
| *envY* | 0.129 | 0.021 | 0.1537 |
| *phoB* | 0.129 | 0.015 | 0.1537 |
| *uhpA* | 0.129 | 0.012 | 0.1537 |
| *fucR* | 0.129 | 0.007 | 0.1543 |
| *malT* | 0.129 | 0.002 | 0.1543 |
| *hdfR* | 0.129 | 0.011 | 0.1550 |
| *pdhR* | 0.130 | 0.014 | 0.1556 |
| *gcvA* | 0.130 | 0.011 | 0.1563 |
| *zraR* | 0.131 | 0.015 | 0.1582 |
| *trpR* | 0.133 | 0.013 | 0.1614 |
| *cusR* | 0.135 | 0.028 | 0.1659 |

| | | | |
|---|---|---|---|
| *hyfR* | 0.135 | 0.005 | 0.1659 |
| *baeR* | 0.135 | 0.006 | 0.1659 |
| *deoR* | 0.135 | 0.009 | 0.1665 |
| *yqhC* | 0.136 | 0.015 | 0.1672 |
| *pepA* | 0.136 | 0.015 | 0.1678 |
| *ompR* | 0.136 | 0.045 | 0.1684 |
| *yiaJ* | 0.136 | 0.017 | 0.1684 |
| *tdcR* | 0.137 | 0.009 | 0.1704 |
| *yjiE* | 0.138 | 0.016 | 0.1717 |
| *cpxR* | 0.139 | 0.013 | 0.1742 |
| *hipB* | 0.139 | 0.010 | 0.1742 |
| *ascG* | 0.140 | 0.009 | 0.1755 |
| *putA* | 0.140 | 0.016 | 0.1761 |
| *dinJ* | 0.142 | 0.022 | 0.1800 |
| *qseB* | 0.144 | 0.023 | 0.1832 |
| *agaR* | 0.144 | 0.008 | 0.1838 |
| *trpD* | 0.145 | 0.014 | 0.1845 |
| *trpE* | 0.146 | 0.011 | 0.1864 |
| *iclR* | 0.146 | 0.007 | 0.1870 |
| *dhaR* | 0.150 | 0.015 | 0.1954 |
| *cysB* | 0.155 | 0.001 | 0.2043 |
| *ihfA* | 0.158 | 0.024 | 0.2095 |
| *hns* | 0.160 | 0.008 | 0.2146 |
| *flhC* | 0.166 | 0.018 | 0.2255 |
| *trpB* | 0.168 | 0.017 | 0.2287 |
| *yqjI* | 0.175 | 0.022 | 0.2428 |
| Standard Curve | | | |
| 0 mmol | 0.047 | 0.004 | |
| 0.2 mmol | 0.184 | 0.002 | |
| 0.4 mmol | 0.256 | 0.049 | |
| 0.6 mmol | 0.358 | 0.033 | |
| 0.8 mmol | 0.473 | 0.044 | |
| 1 mmol | 0.522 | 0.028 | |
| 1.2 mmol | 0.560 | 0.052 | |

[1]Mean absorbance of 3 biological replicates taken at 540 nm.

**Supplementaty Table 2. Total indole concentrations of *E. coli* transcription factor deletants in M9 minim**

| *Gene* | Mean absorbance[1] | STDEV | mmols of Indole |
|---|---|---|---|
| **WT** | 0.039 | 0.002 | 0.0000 |
| *fnr* | 0.043 | 0.000 | 0.0000 |
| *soxS* | 0.039 | 0.002 | 0.0000 |
| *crp* | 0.040 | 0.001 | 0.0000 |
| *lysR* | 0.040 | 0.002 | 0.0000 |
| *fucR* | 0.039 | 0.002 | 0.0000 |
| *malI* | 0.040 | 0.001 | 0.0000 |
| *phoB* | 0.039 | 0.002 | 0.0000 |

| | | | |
|---|---|---|---|
| *cpxR* | 0.039 | 0.002 | 0.0000 |
| *creB* | 0.038 | 0.001 | 0.0000 |
| *trpB (No growth)* | 0.000 | 0.000 | 0.0000 |
| *trpD No growth)* | 0.000 | 0.000 | 0.0000 |
| *trpE No growth)* | 0.000 | 0.000 | 0.0000 |
| *paaX* | 0.039 | 0.001 | 0.0000 |
| *trpA No growth)* | 0.000 | 0.000 | 0.0000 |
| *tnaA* | 0.038 | 0.001 | 0.0000 |
| *trpL* | 0.039 | 0.002 | 0.0000 |
| *tnaC* | 0.040 | 0.002 | 0.0000 |
| *tnaB* | 0.040 | 0.002 | 0.0000 |
| *dhaR* | 0.040 | 0.001 | 0.0000 |
| Standard Curve | | | |
| 0 mmol | 0.043 | 0.000 | |
| 0.2 mmol | 0.284 | 0.019 | |
| 0.4 mmol | 0.403 | 0.037 | |
| 0.6 mmol | 0.454 | 0.003 | |
| 0.8 mmol | 0.480 | 0.040 | |
| 1 mmol | 0.507 | 0.053 | |
| 1.2 mmol | 0.502 | 0.060 | |

[1]Mean absorbance of 3 biological replicates taken at 540 nm.
